# Direct-To-Consumer DNA testing of 6,000 dogs reveals 98.6-kb duplication causing blue eyes and heterochromia in Siberian Huskies

**DOI:** 10.1101/247973

**Authors:** P. E. Deane-Coe, E. T. Chu, A. R. Boyko, A. J. Sams

## Abstract

Consumer genomics enables genetic discovery on an unprecedented scale by linking very large databases of personal genomic data with phenotype information voluntarily submitted via web-based surveys^1^. These databases are having a transformative effect on human genomic research, yielding insights on increasingly complex traits, behaviors, and disease by including many thousands of individuals in genome-wide association studies (GWAS)^2, 3^. The promise of consumer genomic data is not limited to human research, however. Genomic tools for dogs are readily available, with hundreds of causal Mendelian variants already characterized^4–6^, because selection and breeding have led to dramatic phenotypic diversity underlain by a simple genetic structure^7, 8^. Here, we report the results of the first consumer genomics study ever conducted in a non-human model: a GWAS of blue eyes based on more than 3,000 customer dogs with a validation panel of nearly 3,000 more, the largest canine GWAS to date. We discovered a novel association with blue eyes on chromosome 18 (*P* = 1x10^-65^) and used both sequence coverage and microarray probe intensity data to identify the putative causal variant: a 98.6-kb duplication directly upstream of the hox gene *ALX4*, which plays an important role in mammalian eye development^9, 10^. This duplication was largely restricted to Siberian Huskies and is highly, but not completely, penetrant. These results underscore the power of consumer-data-driven discovery in nonhuman species, especially dogs, where there is intense owner interest in the personal genomic information of their pets, a high level of engagement with web-based surveys, and an underlying genetic architecture ideal for mapping studies.

## Main Text

Humans have been exerting multifarious selection on dogs since their domestication from wolves, including strong natural selection during adaptation to a domesticated lifestyle followed by intense artificial selection during modern breed formation^11–13^. One unintended consequence of this selection is that the canine genome now encodes dramatic phenotypic diversity highly amenable for genetic mapping, with moderate genome-wide divergence between breeds except near loci under selection^4, 7, 8^ and long tracts of linkage disequilibrium that can be effectively scanned with microarrays^14^. Genetic discoveries in dogs benefit breeding efforts and animal welfare, and they are valuable for translational studies in humans because dogs and humans exhibit many analogous physical traits, behaviors, and diseases in a shared environment^7, 15^.

In humans, blue eyes first arose in Europeans^16^ and may have been favored by sexual selection due to an aesthetic preference for rare phenotypic variants^17^, as an informative recessive marker of paternity^18^, and/or as a by-product of selection for skin de-pigmentation to increase UVB absorption^19^. Whatever the cause, this selection has acted on the regulatory machinery of *OCA2* (Oculocutaneous Albinism II Melanosomal Transmembrane Protein), which controls transport of the melanin precursor tyrosine within the iris^20, 21^. Because blue eyes result from reduced melanin synthesis, other mutations affecting melanocyte function can also recapitulate the phenotype^22^.

In dogs, blue eyes are iconic of the Siberian Husky, a breed of northern latitudes. Prized among breeders, it is not known whether blue eyes confer adaptive benefits for high latitude dogs as has been hypothesized for humans, and the genetic basis has not yet been discovered. According to breeders, blue eyes in Siberian Huskies are a common and dominant trait, including solid blue and complete heterochromatism (one blue and one brown eye), whereas blue eyes appear to be a rare and recessive trait in breeds like the Border Collie, Pembroke Welsh Corgi, and Old English Sheepdog. The only genetic factors known to produce blue eyes are two cases associated with coat coloration: “Merle” and “piebald” dogs have patchy coat colors due to mutations in Premelanosome Protein *(PMEL17)* and Melanogenesis Associated Transcription Factor *(MITF)* that can lead to one or two blue eyes when de-pigmented regions extend across the face^23, 24^.

We employed a novel genomic resource—a panel of 6,092 dogs genetically tested on a high-density 214,661-marker platform, with owners that had contributed phenotype data via web-based surveys—to examine the genetics of blue eyes in a diverse panel of purebred and mixed-breed dogs.

Using a discovery panel of 3,180 dogs, we detected two significant associations with blue eyes, one on chromosome 10 at *PMEL17* (“merle”; canFam3.1 position 292,851; *P* = 7x10^-49^) and a novel locus on chromosome 18 (CFA18) that had not been previously characterized (position 44,924,848; *P* = 1x10^-68^; Figures 1A, S1). Markers near *MITF* were not significantly associated with blue eyes (*P* = 0.02–0.90 from positions 21,834,567–21,848,176 on CFA20), likely because piebald coat color causes blue eyes in only a small subset of cases.

**Figure 1.**
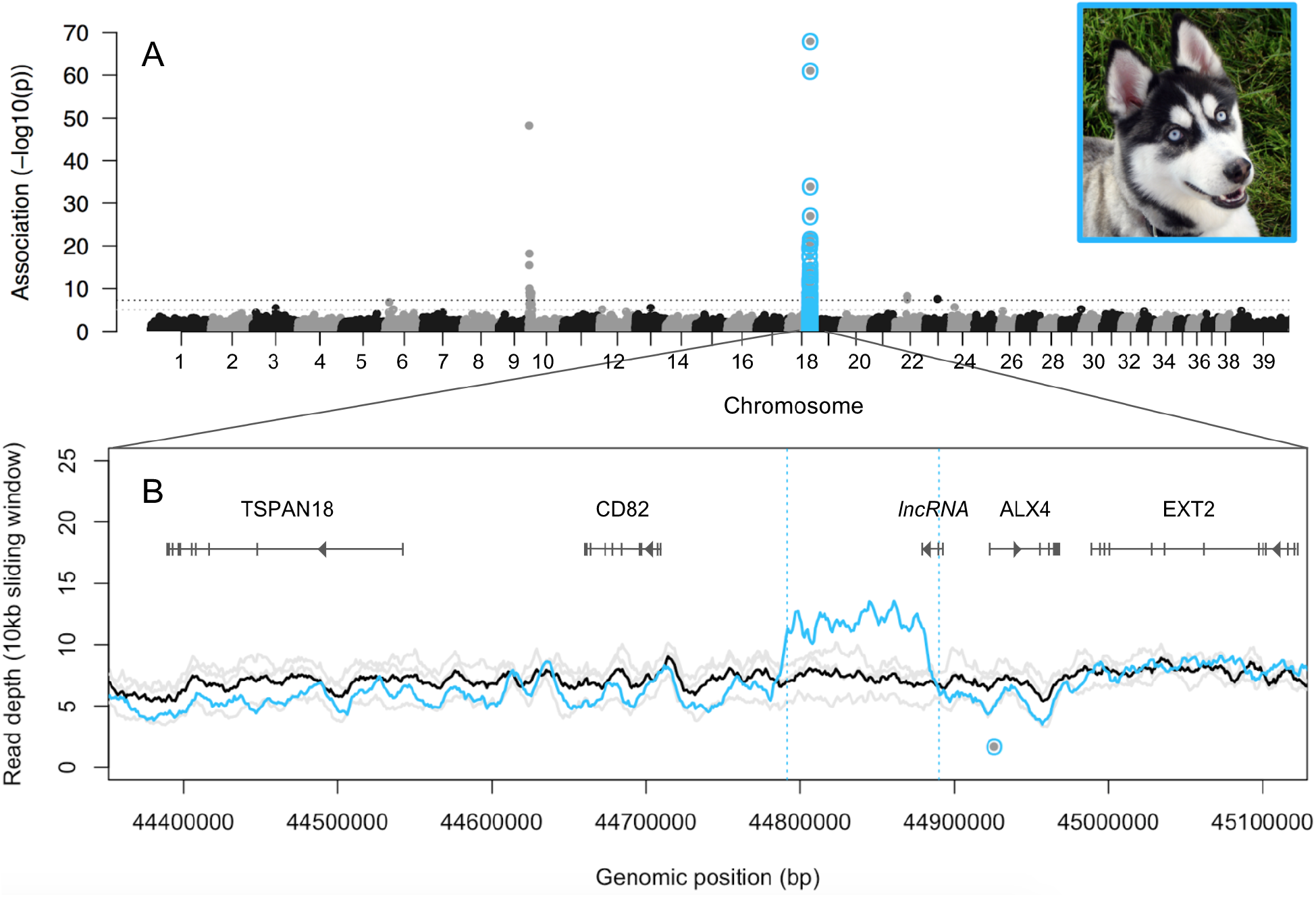
A) Manhattan plot of associations with blue vs. brown eyes across the genomes of 3,180 dogs. Horizontal lines represent the thresholds for suggestive (grey; *P* < 1x10^-5^) and significant (black; *P* < 5x10^-8^) associations. B) Read depth in 10-kb sliding windows across the CFA18 GWAS peak region, for three dogs (grey=individual tracks; black=mean) who do not carry the allele associated with blue eyes (dot at 44,924,848), and one Siberian Husky who does (blue). Blue vertical lines indicate paired-end reads that aligned 98.6-kb from their mate and in an opposite orientation. Photo credit: Pamela Carls.

The novel association on CFA18 was robust to whether heterochromia (complete or sectoral) was considered (solid blue only *P* = 3x10^-71^, heterochromia only *P* = 1x10^-12^; Figure S2), and remained strong when we restricted our analysis to only purebred or mixed-breed dogs (purebred *P* = 3x10^-9^, mixed-breed *P* = 3x10^-63^; Figure S3). Although the minor allele (A) at the CFA18 locus was carried (in one or two copies) by only 10% of dogs in this dataset (both blue-and brown-eyed), it was carried by 78% of non-merle blue-eyed dogs (32% homozygous, 68% heterozygous) and 100% of blue-eyed purebred Siberian Huskies (*N* = 22).

We defined a fine-mapping panel using the 314 dogs from the discovery panel that did not carry merle, carried at least one copy of the CFA18 allele associated with blue eyes, and for which supplemental Illumina microarray data were available. Of these, 87 (26%) had at least one blue eye. Blue-eyed dogs homozygous for the CFA18 marker (*N* = 26) exhibited a long shared haplotype block in the region containing that SNP (Figure S4); however, we observed four SNPs within the block (positions 44800358, 44822014, 44825760 and 44849276) that were consistently heterozygous, suggesting a nonbalanced structural variant overlapping those markers in dogs carrying the blue-eyed haplotype.

Canine whole genome sequences available on the NCBI Sequence Read Archive (SRA) included one Siberian Husky heterozygous for the CFA18 allele associated with blue eyes. Genome-wide read depth for the Siberian Husky was 8x, but coverage abruptly increased to 12x (a 1.5x increase) across an intergenic region from 44.79–44.89-Mb that encompassed the four heterozygous SNPs (Figure 1B). Furthermore, 30% of the paired-end reads spanning 44,791,417–44,791,584 had a mate that mapped in an opposite orientation to positions 44,890,024–44,890,166, consistent with a 98.6-kb tandem duplication for which the midpoint span was less than the insert size of the paired end reads (< 350-bp)^25, 26^. This pattern was not observed in sequenced dogs that did not carry the CFA18 minor allele.

We compared log-transformed intensity data (*log R*) for SNPs inside vs. outside the duplication (Δ *log R*) in our fine-mapping panel and observed that blue-eyed dogs exhibited higher values of Δ *log R* (0.15–0.56) than brown-eyed dogs (range; *P* < 2x10^-16^), consistent with the hypothesis that blue-eyed dogs carry one or two additional copies of probe sequences for SNPs within the duplicated region (Figure 2A). Five outlier dogs did not appear to carry the duplication despite being blue-eyed; however, most of these had atypical coat pigmentation (Supplemental Information). Blue-eyed heterozygotes and homozygotes for the duplication (*N* = 82 / 314) exhibited different distributions in Δ *log R* (*P* = 4x10^-12^; Figure 2B). The high proportion of blue-eyed heterozygotes suggests that the duplication is dominant in its phenotypic effect, however the bimodal distribution of Δ *log R* among brown-eyed dogs suggests that one or more other factors may modify or override the duplication’s effect on eye color (Figure 2C; Supplemental Information).

**Figure 2.**
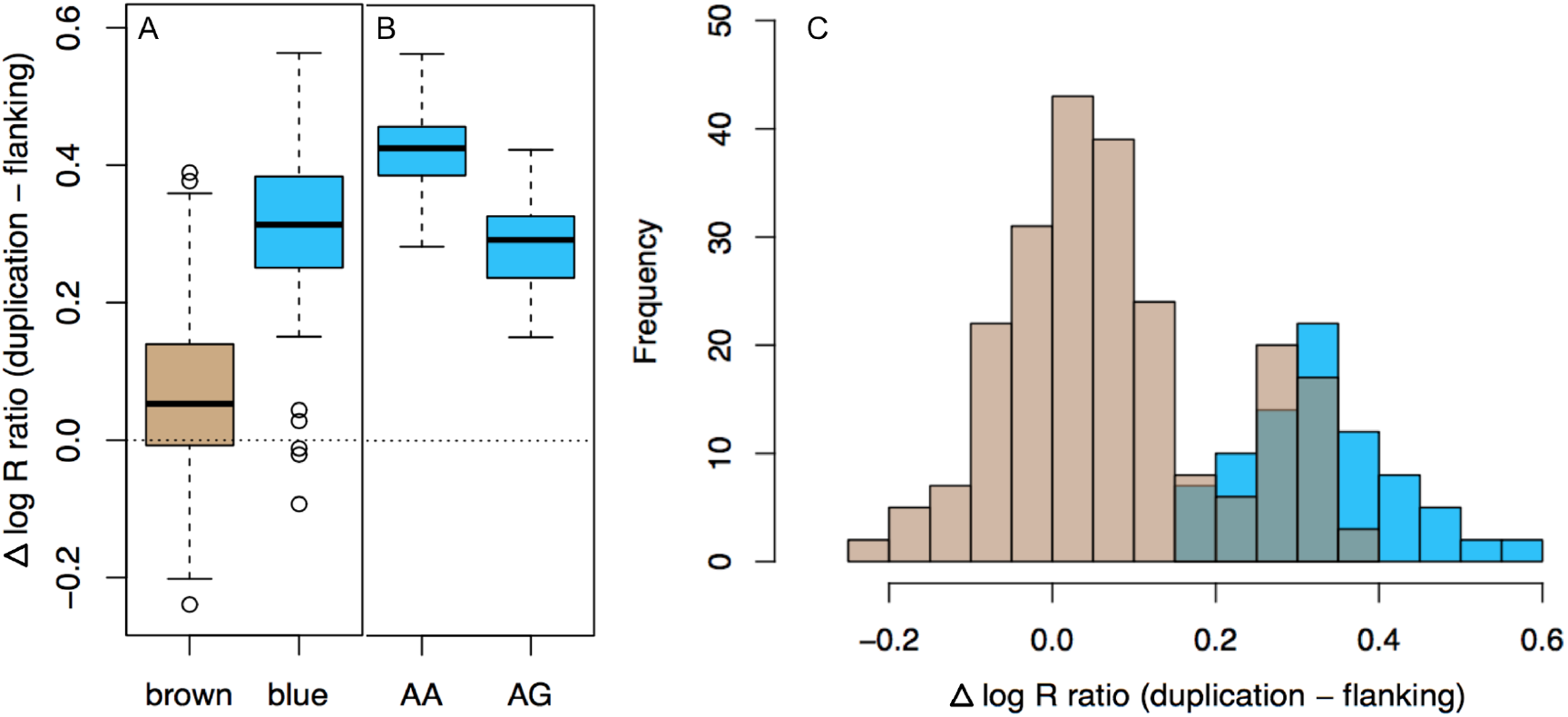
A) Among non-merle carriers of the CFA18 GWAS allele, blue-eyed dogs exhibited higher *log R* intensity at SNPs within the duplication than in flanking regions (Δ *log R* = 0.15–0.56, excluding five outliers) compared to brown-eyed dogs (*P* < 2.2 x 10^-16^; *N* = 314). B) Blue-eyed dogs heterozygous vs. homozygous for the CFA18 allele exhibited distinct distributions in Δ *log R* (*P* = 4.0 x 10^-12^; *N* = 82), consistent with being heterozygous vs. homozygous for the duplication. C) Some brown-eyed dogs had Δ *log R* values within the range of blue-eyed duplication carriers (*N* = 35 / 309).

We compiled a database of 2,912 purebred customer dogs distinct from those included in our GWAS panel to quantify the prevalence of the duplication in different breeds and to perform a validation test of its association with blue eyes. We defined the first quartile of the Δ *log R* distribution among blue-eyed GWAS dogs (Δ *log R* > 0.27) as a conservative minimum threshold for detecting duplication carriers. Using this threshold 2% of dogs in the validation panel carried the duplication (*N* = 63 / 2,912), and 90% of these were Siberian Huskies. The remainder were Klee Kai, a breed derived from Siberian Husky (*N* = 2), Australian Shepherd (*N* = 3), and one German Shepherd. Profile photos were available for 75% of dogs with the duplication (*N* = 49 / 63), and all but three had blue or heterochromatic eyes instead of solid brown. Two Siberian Huskies and the German Shepherd had brown eyes despite having Δ *log R* values consistent with being heterozygous for the duplication (0.31–0.32). The owners of two of these dogs were able to provide additional information that confirmed they were likely carriers: One of the huskies had blue-eyed parents and sired all blue-eyed or heterochromatic litters, and the German Shepherd had a heterochromatic sire and littermates.

To date, the most familiar examples of duplications affecting phenotype are those related to dosage, cases where one or more duplication events increased gene copy number and, therefore, the amount of translated protein product available for cellular function^27, 28^. However, this duplication sits in an intergenic region between the tetraspanin *CD82* and hox gene *ALX4* (NCBI; UCSC Genome Browser). Two non-coding RNAs (ncRNAs) are annotated on the complementary strand, including an uncharacterized long noncoding RNA (lncRNA) that overlaps the 3’ breakpoint of the duplication (Figure 1; Figure S4–S5). We could find no evidence that *CD82* is functionally associated with eye color in humans or any other animal, but *ALX4* and its paralogs play an important role in mammalian eye development^9, 10^. Given the importance of *cis*-regulatory elements in local gene regulation^29, 30^, we propose that this duplication could cause blue eyes by disrupting regulation of *ALX4.*

Functional follow-up studies are needed to explicitly assay regulatory changes in *ALX4* caused by this duplication; however, we have shown that this mutation is highly (but not completely) penetrant and largely restricted to Siberian Huskies. Of the 62 blue-eyed dogs in our discovery panel without merle or piebald (out of 156 blue-eyed dogs), this duplication explains the blue-eyed phenotype in a majority of them (33 dogs carry the duplication using the conservative Δ log R > 0.27 detection threshold). By using consumer genomic data to drive this research, we were able to build the largest canine GWAS dataset to date, determine the prevalence of the putative causal variant across a diverse population, and recontact owners of specific dogs to learn more about the inheritance of the trait. As more canine genetic testing is done on high-density array platforms, these databases hold particular promise for unlocking the genetic basis of complex phenotypes for which dogs are a particularly useful model, including cancer, behavior, and aging.

## Data availability

Additional data and complete summary statistics from the analyses in this paper will be made available to researchers through Embark Veterinary, Inc, under an agreement with Embark that protects the privacy of Embark customers and their dogs. Please contact the corresponding author for more information and to apply for access to the data.

## Acknowledgements

We thank customers of Embark who participated in online surveys on behalf of their dogs, as well as all the Embark employees who made this work possible, particularly Ryan Boyko, Matt Barton, Adam Gardner, Tiffany Ho and David Riccardi. We also thank our scientific advisors, Cornell University, and the Kevin M. McGovern Family Center for Venture Development in the Life Sciences, for their guidance and encouragement. This study was funded by the participants and by Embark.

## Author Contributions

PED analyzed the data and wrote the paper. ARB and AJS jointly directed the research and writing. ETC conducted additional analyses and background research, and contributed edits to the paper.

## Author Information

Conflict of interest statement: PED, ETC, ARB and AJS are employees of Embark Veterinary, a canine DNA testing company which will offer commercial testing for the variant described in this study. ARB is co-founder and part owner of Embark. Correspondence and requests for materials should be addressed to PED, ARB, or AJS.

## Methods

### Discovery samples

We solicited phenotype data from customers whose dogs have been genetically tested by Embark Veterinary, and who have agreed to participate in research, by implementing an online survey about their dog’s morphological traits at http://embarkvet.com and encouraging participation via email. We initiated the survey on February 7, 2017, and, as of November 23, 2017, owners of 3,248 adolescent and adult dogs whose eye color can be assumed to be developmentally complete (6 months or older) had submitted a response to the section of that survey that asks about eye color (Figure S6). A subset of these owners (*N* = 68) selected “other”, indicating that their dog had an eye color not represented by any of the seven options. In total, 156 dogs in this dataset were reported to have either solid blue eyes (*N* = 73) or heterochromic eyes (partially blue; *N* = 83), compared to 3,024 with some shade of solid brown. We encoded this trait as a binary phenotype in case-control format (0: brown, 1: blue) and considered both solid blue and heterochromic dogs as cases. In total, 21% of these dogs were purebreds of various breeds, and the remainder were mixes of two or more breeds. Ancestry from 185 different dog breeds or landraces was represented in this dataset.

### Genotyping & Quality Control

Customer dogs were genotyped on Embark’s custom high-density 214,661-marker platform, ensuring genotype concordance rates above 99.99% and missingness rates below 0.1%. Total genotyping rate was 99.5%, and all dogs (N = 3,180) had less than 2.5% missing data and passed standard filtering in PLINK^31^. After filtering, 89.7% of variant sites (192,550 / 214,661) were genotyped across at least 95% of individuals and were included in subsequent analyses.

### Genome-wide association

We constructed a relatedness matrix from centered genotypes and ran a genome-wide association test based on a univariate linear mixed-model in GEMMA^32^, using the eigenvalues and eigenvectors of the relatedness matrix to control for confounding effects of shared ancestry, particularly among dogs of the same breed or breed group (groups of closely related or recently derived breeds). We identified significant associations by applying a threshold of *P* < 5x10^-8^ to the Wald test statistic.

### Whole genome sequence analysis

We downloaded whole genome sequence data for 325 different dog breeds from the NCBI Sequence Read Archive (www.ncbi.nlm.nih.gov/sra), calculated read depth coverage across sites using SAMtools^33^, and investigated mapped paired-end reads in regions of interest using the Integrative Genomics Viewer (IGV)^34, 35^.

### Validation samples

The validation panel included purebred representatives from 194 different breeds, eight village dog landraces, and gray wolves. We assayed the ratio of log-transformed intensity for samples labeled with red vs. green dye on the microarray (*log R* ratio) for 2,769 phenotyped non-merle dogs in our database at 50 markers, eight within the putative bounds of the duplication (positions 44800358, 44822014, 44825760, 44838433, 44849276, 44855038, 44858831, 44876627), and 42 markers located directly upstream and downstream. We compared the change in intensity between duplication markers and flanking regions (Δ *log R*) according to both the eye color phenotype and GWAS SNP genotype of each dog.

### Data availability

Embark will make complete summary statistics from this dataset available to researchers under an agreement that protects the privacy of the Embark customers and their pets. Please contact the corresponding author for more information and to apply for access to the data.

## Supplemental Information

*Other blue eye phenotypes*. Of the dogs in our discovery panel with blue eyes not explained by either merle (*N* = 92 / 156) or the CFA18 marker (*N* = 41 / 156), 35% were blue-eyed due to white facial markings (e.g. piebald) according to profile photos uploaded by their owners (*N* = 8). The remainder were merle cases not predicted by the CFA10 merle-associated SNP (*N* = 4), other eye colors misreported as blue (*N* = 3), or were unknown (N = 8 without profile photos). Of the five outlier dogs that did not appear to carry the duplication, despite being blue-eyed and carrying the CFA18 marker, four had atypical coat pigmentation (leucistic, mostly white, or unusual piebald-like cases). The fifth is a mixed breed dog with unexplained sectoral heterochromia.

### Supplemental Figures

**Figure S1.**
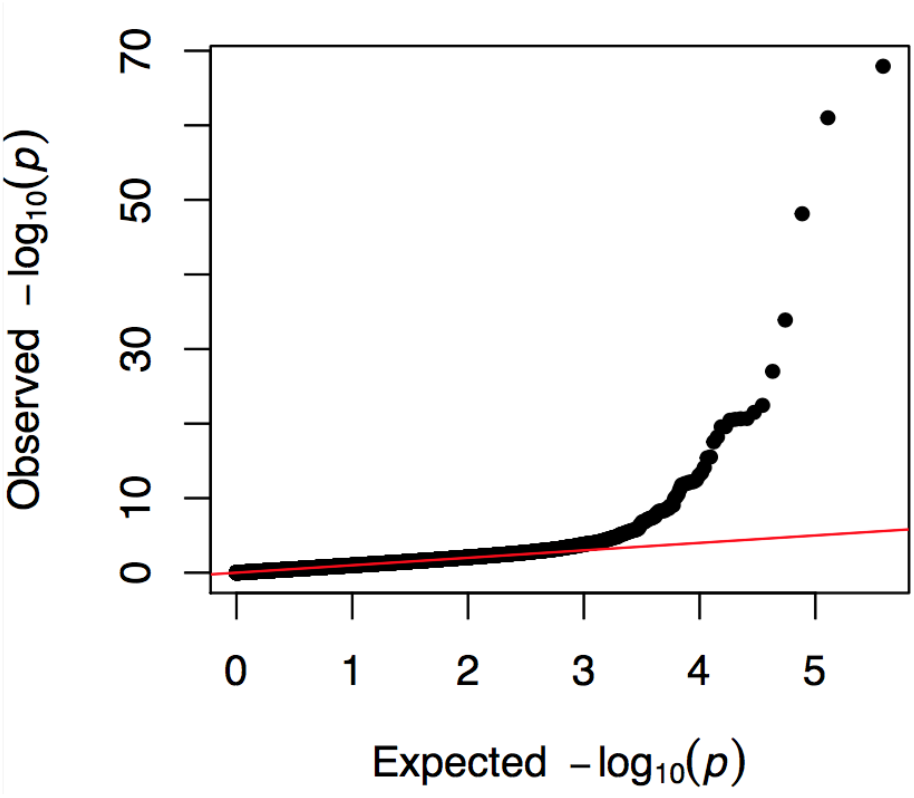
QQ plot of the association between genotype and blue vs. brown eyes across the genomes of 3,180 purebred and mixed-breed dogs (corresponds to Manhattan plot in Figure 1).

**Figure S2.**
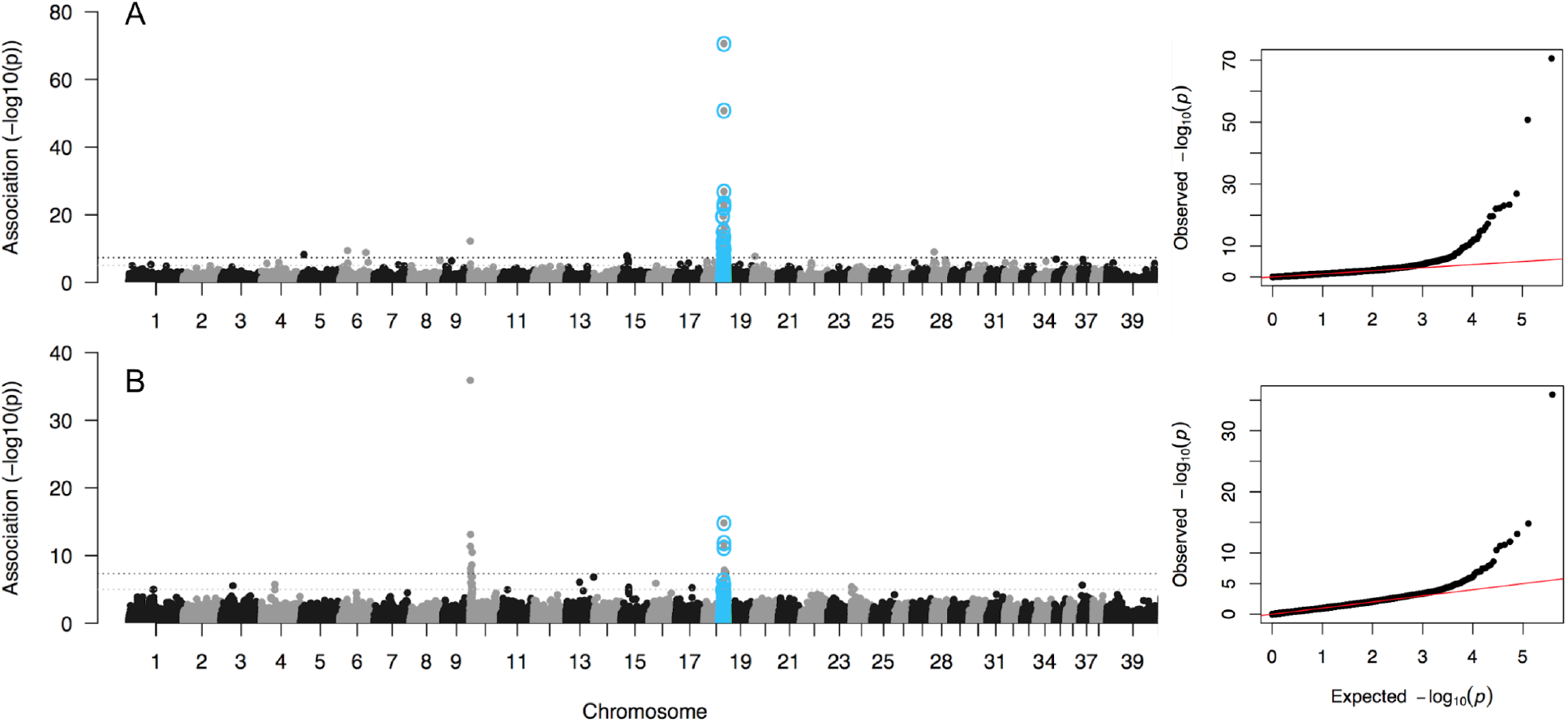
Manhattan and QQ plots of genome-wide associations with A) solid blue eyes (73 cases) and B) heterochromia (83 cases) across the genomes of 3,180 purebred and mixed breed dogs (192,550 markers). Grey and black dotted horizontal lines represent the thresholds for suggestive (*P* < 1x10^-5^) and significant (*P* < 5x10^-8^) associations, respectively.

**Figure S3.**
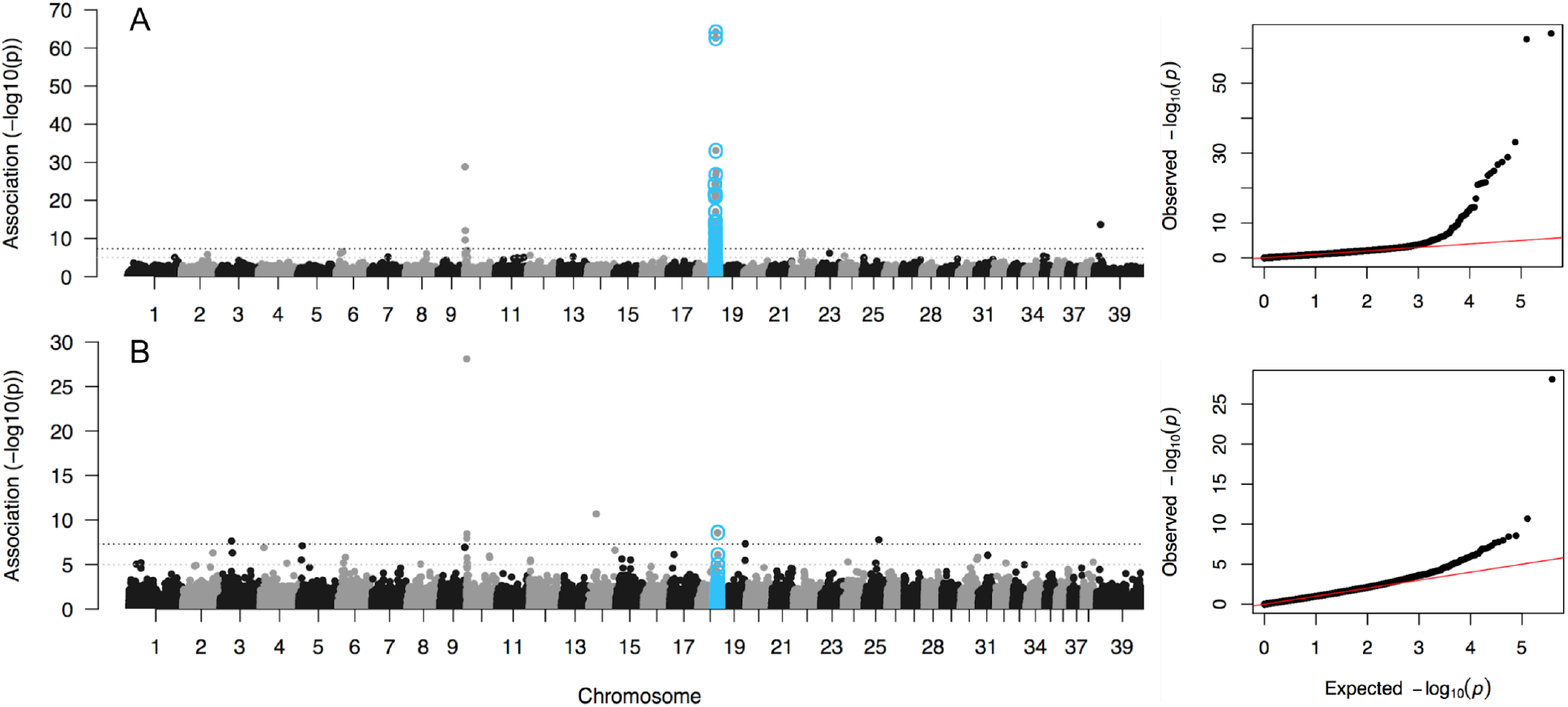
Manhattan and QQ plots of genome-wide associations for A) 2,448 mixed-breed dogs (192,570 markers) and B) 670 purebred dogs (191,854 markers). Grey and black dotted horizontal lines represent the thresholds for suggestive (*P* < 1x10^-5^) and significant (*P* < 5x10^-8^) associations, respectively.

**Figure S4.**
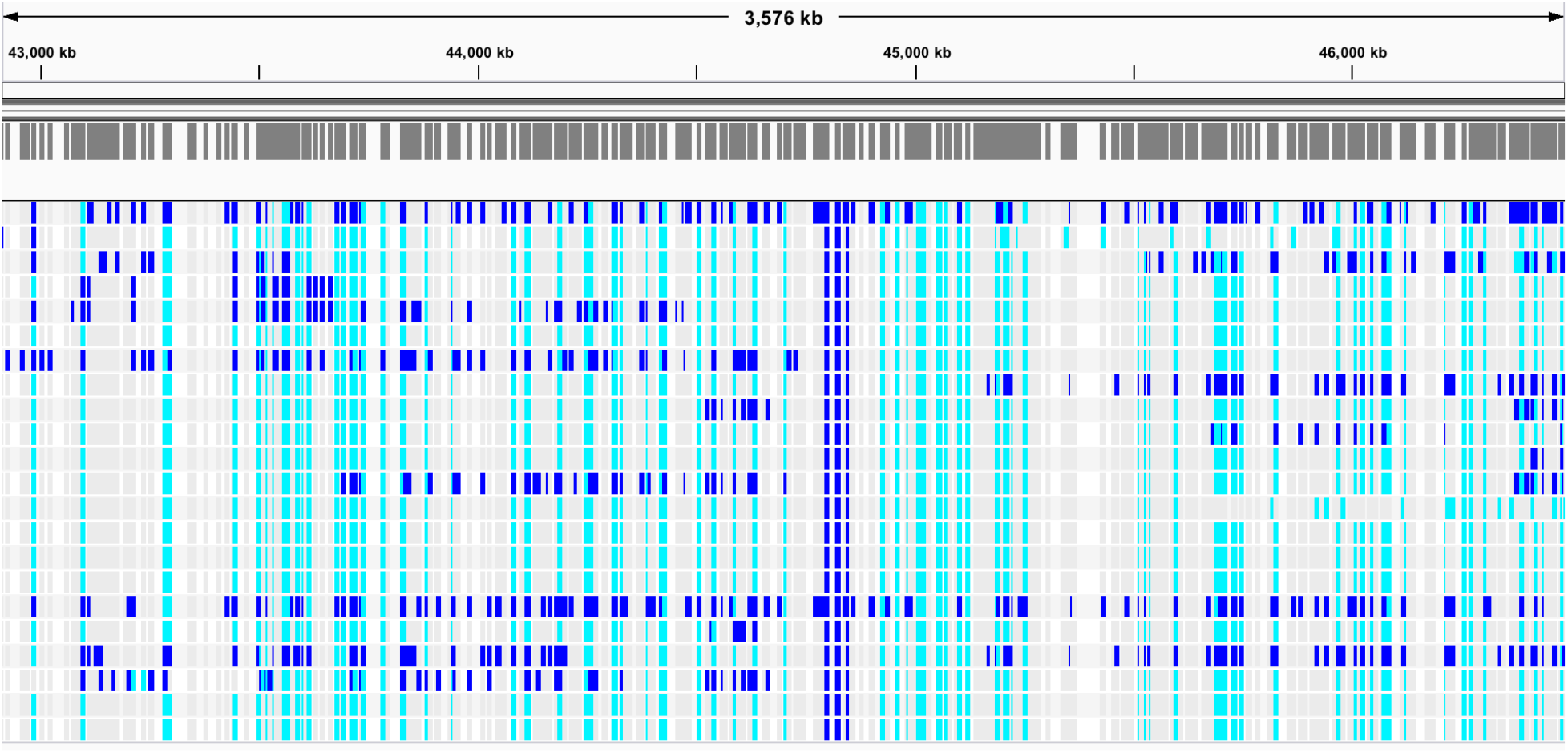
Bounds of a linked haplotype block present in dogs homozygous for the CFA18 allele associated with blue eyes, featuring four heterozygous markers (positions 44,800,358–44,849,276) suggestive of a non-balanced structural variant (teal = homozygous, blue = heterozygous).

**Figure S5.**
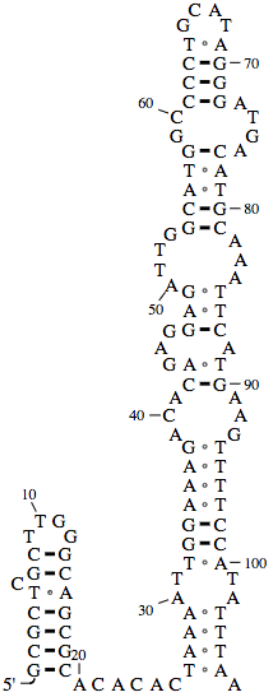
Structural diagram for snRNA located within duplicated region (UCSC Genome Browser).

**Figure S6.**
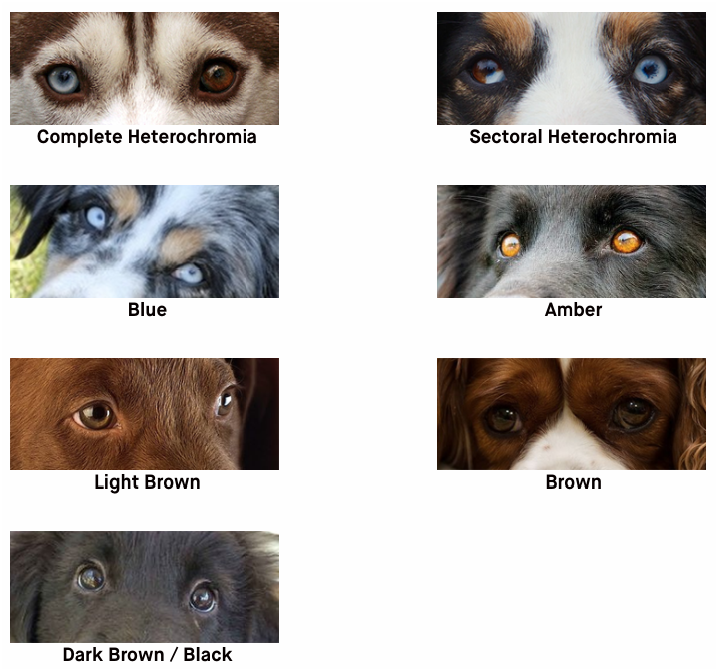
Participants were prompted to report their dog’s eye color as one of seven options, guided by visual examples (images courtesy of musingsofabiologistanddoglover.blogspot.com and quaggatale.wordpress.com).

